# Merkel Cell Polyomavirus Small Tumor Antigen Activates Matrix Metallopeptidase-9 Gene Expression for Cell Migration and Invasion

**DOI:** 10.1101/2020.03.02.974303

**Authors:** Nnenna Nwogu, Luz E. Ortiz, Adrian Whitehouse, Hyun Jin Kwun

**Affiliations:** Department of Microbiology and Immunology, Penn State University College of Medicine, Hershey, PA, USA; Penn State Cancer Institute, Hershey, PA, USA; School of Molecular and Cellular Biology and Astbury Centre for Structural Molecular Biology, Faculty of Biological Sciences, University of Leeds, Leeds, United Kingdom

**Author notes:** Correspondence: Hyun Jin Kwun, Department of Microbiology and Immunology, Penn State University College of Medicine, Hershey, PA 17033, USA, E-mail addresses. Competing interests: The authors declare no competing interests.

**Keywords:** Merkel cell carcinoma, Merkel cell polyomavirus, small tumor antigen, Matrix Metallopeptidase-9, FBW7, Cell migration, Cell invasion

## Abstract

Merkel cell polyomavirus (MCV) small T antigen (sT) is the main oncoprotein for the development of Merkel cell carcinoma (MCC). MCC is a rare, clinically aggressive neuroendocrine tumor of the skin with a high propensity for local, regional, and distant spread. The dysregulation of matrix metalloproteinase-9 (MMP-9) has been implicated in multiple essential roles in the development of various malignant tumor cell invasion and metastasis. Previously, MCV sT was shown to induce the migratory and invasive phenotype of MCC cells through the transcriptional activation of the Sheddase molecule, ADAM 10 (A disintegrin and metalloprotease domain-containing protein 10). In this study, we show that MCV sT protein stimulates differential expression of epithelial–mesenchymal transition (EMT) associated genes, including MMP-9 and Snail. This effect is dependent on the presence of the large T stabilization domain (LSD), which is known to be responsible for cell transformation through targeting of promiscuous E3 ligases, including FBW7, a known MMP-9 and Snail regulator. Chemical treatments of MMP-9 markedly inhibited sT-induced cell migration and invasion. These results suggest that MCV sT contributes to the activation of MMP-9 as a result of FBW7 targeting, and increases the invasive potential of cells, which can be used for targeted therapeutic intervention.

**IMPORTANCE:** Merkel cell carcinoma (MCC) is the most aggressive cutaneous tumor without clearly defined treatment. Although MCC has a high propensity for metastasis, little is known about the underlying mechanisms that drive MCC invasion and metastatic progression. MMP-9 has shown to play a detrimental role in many metastatic human cancers, including melanoma and other non-melanoma skin cancers. Our study shows that MCV sT-mediated MMP-9 activation is driven through the LSD, a known E3 ligase targeting domain, in MCC. MMP-9 may serve as the biochemical culprit to target and develop a novel approach for the treatment of metastatic MCC.

## INTRODUCTION

Merkel cell carcinoma (MCC) is a rare skin cancer of neuroendocrine origin with a high propensity to metastasize (1). Although the incidence rate of MCC is lower than melanoma, it is highly aggressive with an estimated mortality rate of 33%-46*%*; hence it is significantly more lethal than malignant melanoma (2). Merkel cell polyomavirus (MCV) is the etiological agent of MCC. The majority of MCC cases are associated with MCV as observed by monoclonal integration of the MCV genome in the tumor DNA (3). As a classic polyomavirus, the genomic organization of MCV is similar to other known human polyomaviruses. MCV expresses small and large tumor antigens (sT and LT), which are essential for viral replication and pathogenesis. MCC tumor-derived MCV LT sequences integrated into MCC genomes contain mutations prematurely truncating the C-terminal growth inhibitory domain (4), while MCV sT remains intact.

MCV sT has been shown to mediate multiple oncogenic mechanisms that contribute to MCC development. Inhibition of SCF (Skp1, Cullin, F-box containing complex) E3 ligases by MCV sT appears to induce several viral and cellular oncoprotein activation, leading to enhanced MCV replication and cell proliferation (5, 6). Aberrant activation of oncogenic potential in MCV sT expressing cells also promoted the malignant phenotypes that are involved in genomic instability such as centrosome amplification, aneuploidy, and micronuclei formation (7). This oncogenic activity of MCV sT requires the LT stabilization domain (LSD), a unique and disordered domain of MCV sT, which is known to interact with SCF E3 ligase complexes (5, 7). Although the exact mechanism by which MCV sT targets E3 ligases is yet to be elucidated, it is clear that the LSD plays a significant role in the distinctive transforming activities induced by MCV sT in vitro and in vivo (5–7).

The E3 ubiquitin ligase F-box and WD repeat domain containing 7 (FBW7) functions as a putative tumor suppressor and an evolutionarily conserved substrate receptor of SCF ubiquitin ligase complex and plays vital roles in cell proliferation and cell migration (8). In various cancers, including renal cancer (9, 10), gastric cancer (11) and hepatocellular carcinoma (12), FBW7 inhibition promotes metastasis and epithelial–mesenchymal transition (EMT) by upregulating matrix metalloproteinase expression, specifically MMP-2, MMP-9, and MMP-13. Matrix metalloproteinases (MMPs) are a zinc-dependent family of proteolytic enzymes that participate in the degradation of the extracellular matrix (ECM). Dysregulation of these proteases has been observed in multiple cancers where enhanced expression of certain MMP proteins contribute to cell migration, invasion, and angiogenesis (13, 14). Specifically, MMP-9 has been linked to multiple hallmarks of cancer, including but not limited to metastasis, invasion, immunological surveillance, and angiogenesis (15). MMP-9, also known as 92 kDa type IV collagenase (16), plays a vital role in the degradation of elastin and partially hydrolyzed collagen that is essential for maintaining epithelial structural integrity. Various studies have shown that human tumor virus-associated oncoproteins play a critical role in metastasis and EMT-related mechanisms. Hepatitis B virus (HBV)-encoded X protein (HBx) (17), Kaposi’s sarcoma-associated herpesvirus (KSHV) K1 (18), and Epstein-Barr virus (EBV) latent membrane protein 1 (LMP-1) proteins (19) are known to upregulate MMP-9 expression, thereby contributing to invasiveness and metastasis, key hallmarks of cancer (20).

MCV sT stimulates cell motility by inducing microtubule destabilization (21), actin rearrangement (22) and cell dissociation by disruption of cell junctions (23). Interrogation of previously published quantitative proteomic datasets of MCV sT-expressing cells (21) indicates that MCV sT activated expression of Snail, a transcription factor that enhances mesenchymal genes, and MMP-9. In contrast, MCV sT significantly downregulated genes related to cell adhesion molecules, suggesting the potential function of MCV sT in the regulation of EMT. MMP-9 and Snail activation by MCV sT was strictly dependent on the presence of the LSD, which resulted in the enhancement of cell migration in mouse fibroblast cells and human cancer cell lines. Our findings indicate that MCV sT targeting of cellular E3 ligases may play a role in MCV sT-induced cell migration and invasion in MCC. Notably, chemical treatment with MMP-9 inhibitors resulted in significant inhibition of MCV sT-induced cell migration and invasion. This suggests that MMP-9 protein may be a viable target for novel therapeutic intervention for disseminated MCC.

## RESULTS

### MCV sT expression induces differential expression of proteins associated with EMT

Recent studies have highlighted the involvement of MCV sT in the highly migratory and cell dissociation phenotypes of MCC, elucidating its highly multifunctional roles in MCC (21–23). Previously described SILAC (stable isotope labeling by amino acids in cell culture)-based quantitative proteomics data (21) was further interrogated to assess the alterations in the host cell proteome upon expression of MCV sT in a HEK293 derived cell line (i293-sT) **(Fig. 1A)**(21). These results highlighted an alteration in proteins associated with enhancement of cell migration (microtubule-associated cytoskeletal organization) and cell adhesion as previously reported and the basement membrane proteins, a specialized form of the ECM. Specifically, the quantitative proteomic analysis showed an almost two-fold decrease in Collagen alpha-2(IV) chain (COL4A2) and Laminin subunit gamma 1(LAMC1), two essential components of the basement membrane. The basement membrane is crucial for epithelial structural integrity. It is comprised of a network of glycoproteins and proteoglycans such as Type IV collagen and laminin and provides a barrier from invasion by tumor cells (24). These results suggest that MCV sT plays a role in the basement membrane degradation, an essential process for the metastatic invasion of tumor cells into the circulatory system to occur in MCC. Aligned with these observations, transcriptome analysis has suggested that specific markers associated with EMT are increased upon MCV sT expression (25).

**FIGURE 1.**
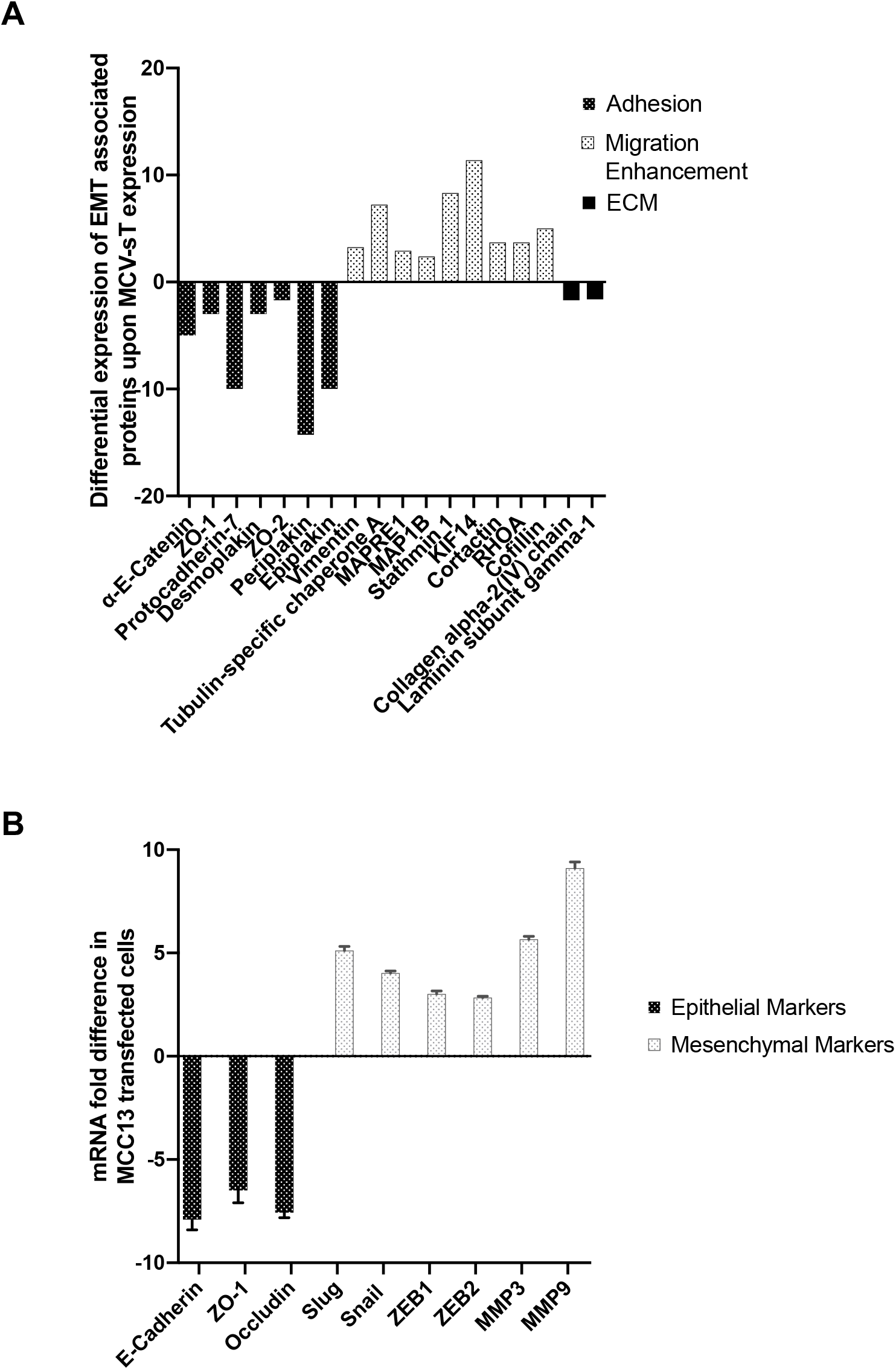
MCV sT leads to differential expression of proteins associated with epithelial to mesenchymal transition (EMT). **(A)** Quantitative proteomics analysis illustrating differential expression of EMT associated proteins upon MCV sT expression. Proteins associated with cell adhesion and structural integrity of the extracellular matrix are downregulated upon MCV sT expression. While expression of proteins which encourage cell migration by reorganization of the actin network and microtubule destabilization are upregulated. **(B)** MCV sT regulates EMT-associated gene expression. MCC13 cells were transfected with control or MCV sT expression plasmids. While epithelial markers were downregulated, mesenchymal markers were significantly upregulated upon MCV sT expression. Cellular RNA was extracted using a trizol reagent and transcript levels were analyzed by RT-qPCR using the comparative ΔΔCt method (n = 3).

To validate the potential regulation of EMT-related markers by MCV sT, RT-qPCR was performed. Changes in mRNA levels of classic EMT markers were assessed in MCC13 cells overexpressing a control of MCV sT construct. Upon MCV sT expression, a significant downregulation in epithelial markers E-cadherin, Zonula occludens-1 (ZO-1) and Occludin was observed **(Fig. 1B)**. Conversely, mesenchymal markers Slug, Snail, ZEB1, ZEB2, MMP-3, and MMP-9 were upregulated upon MCV sT expression. These results infer the possibility of MCV sT inducing an EMT, which contributes to the metastatic potential of MCV-associated MCC.

### MCV sT inhibition of FBW7 contributes to migratory phenotype

An essential requirement of metastasis involves the dissemination of tumor cells to various organs from the primary tumor (26). Multiple studies have demonstrated that loss of FBW7 promotes cell invasion and migration in numerous cancers through modulation of EMT-related cellular factors such as MMP-9 and Snail (27–29), which are upregulated transcriptionally upon MCV sT expression **(Fig. 1B).** The MCV sT LSD region is known to bind and inhibit the FBW7 (5, 7). As a result of this inhibition, FBW7 oncogenic substrates are stabilized in MCV sT-expressing cells, which may contribute to MCV sT-induced migratory phenotype. Although MCV sT and FBW7 interaction has been characterized in vitro by co-immunoprecipitation in over-expressing cell, the in vitro techniques do not identify whether this interaction occurs with endogenous proteins and, therefore, may not reflect the native behavior of their endogenous counterparts. In situ Proximity ligation assay (PLA) can detect interactions with high specificity and sensitivity due to the coupling of antibody recognition and DNA amplification, which provides a technical advantage over other protein-protein interaction assays often plagued with long preparation times and extensive troubleshooting. For that reason, we utilized PLA combined with flow cytometry to revalidate this interaction (30, 31). As shown in **Fig. 2A**, quantification of wild type MCV sT interaction with FBW7 resulted in high-intensity PLA signal comparative to our positive control c-Myc, a well-known FBW7 substrate (32). This interaction was markedly diminished by expression of sT_LSDm_ (**Fig. 2A)**, an LSD alanine mutant of MCV sT, consistent with the finding from a previous report (5).

**FIGURE 2.**
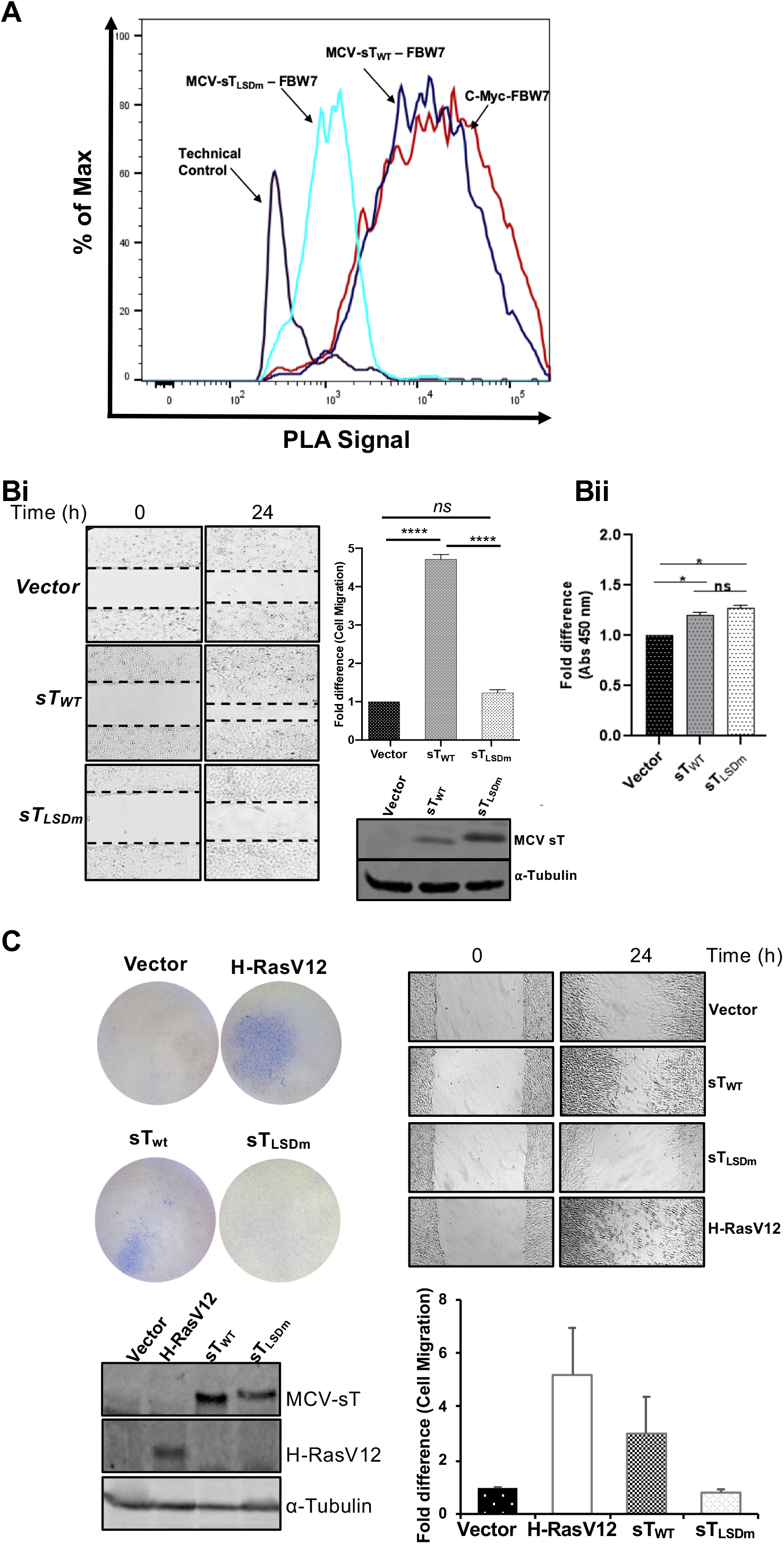
MCV sT induces cell motility in an LSD-dependent manner. **(A)** Validation of MCV sT and FBW7 interaction. To confirm the interaction of HA-FBW7 and MCV sT LSD domain, a PLA-flow cytometric analysis was carried out. Wild type MCV sT displayed an interaction with FBW7 comparative to the positive control interaction of c-Myc with FBW7, while mutation of sT LSD greatly ablated sT interaction with FBW7. Primary antibodies were utilized at optimized concentrations with HA-Tag (C29F4) (1:500), c-Myc (9E10) (1:500), and 2T2 (1:500). Protein expression was evaluated by immunoblot analysis in **Fig. S1A**. **(B)** MCV sT induced-cell migration is LSD-dependent. Scratch assay. Poly-L-lysine-coated 6-well plates were seeded with U2OS cells and transfected with either a vector control, sT_WT_ and sT_LSDm_ (Fbw7 binding mutant) plasmids. Migration of cells toward the scratch was observed over a 24 h period, and images were taken using a REVOLVE4 fluorescent microscope (Echo Laboratories). Scratch assays were performed in triplicate and measured using Fiji Image J analysis software. Differences between means (*p* value) were analyzed using a t-test with GraphPad Prism software. Protein expression was detected by immunoblot analysis to validate successful transfection using 2T2 antibody for sT antigens and α-Tubulin, respectively. No significant differences in cell proliferation were observed between cells expressing MCV sT within 24 h, indicating that cell proliferation does not interfere with the measurement of sT-induced cell migration. **(C)** MCV sT promotes rodent fibroblast cell migration. NIH3T3 cells stably expressing an empty vector, H-RasV12, sT_WT_ and sT_LSDm_ were trypsinized and 2×10^5^ cells were used for transwell migration and scratch assay. H-RasV12 was used as a positive control. The experiments were performed two times, and the results were reproducible. The graph indicates the fold difference of migrated cells relative to the vector control sample. Protein expression was determined by immunoblotting.

To determine whether MCV sT targeting of FBW7 contributes to sT-induced migratory phenotype, a scratch assay was performed comparing vector control, MCV sT_WT_, and MCV sT_LSDm_ in U2OS cells. Images of the scratch area were recorded at time point 0 and 24 h post scratch. Compared to empty vector negative control, MCV sT greatly enhanced the motility and migration of U2OS cells, consistent with previous studies **(Fig. 2B)**(21–23, 33). In contrast, MCV sT_LSDm_ did not show a significant increase in cell migration. Over the 24 h period of the assay, we see no significant positive or negative effect on cell number confirmed by a viability assay, indicating that the resulting phenotype is specific to cell migration **(Fig. 2B)**. Similarly, enhanced cell migration was readily detected with wild type sT in NIH3T3 mouse fibroblast **(Fig. 2C)** and MCC13 (MCV negative MCC cell line) **(Fig. S1B)**, while this phenotype is not induced by sT_LSDm_. This suggests that sT targeting of FBW7 may be involved in the MCV sT-induced cell migratory phenotype.

### MCV sT inhibition of FBW7 prevents turnover of MMP-9

As shown in **Fig. 1B**, MCV sT induces MMP-9, an essential protein associated with the FBW7-EMT axis in human cancers (34). To confirm MCV sT induction of MMP-9 expression, a variety of cell lines, 293, COS-7, MCC13, and U2OS cells were transfected with a vector control and MCV sT plasmids. MMP-9 gene expression is primarily regulated transcriptionally, resulting in low basal levels of these proteases in normal physiology (35). RT-qPCR results showed that MCV sT expression significantly increased MMP-9 transcript levels in all cell lines tested (**Fig. 3A**).

**FIGURE 3.**
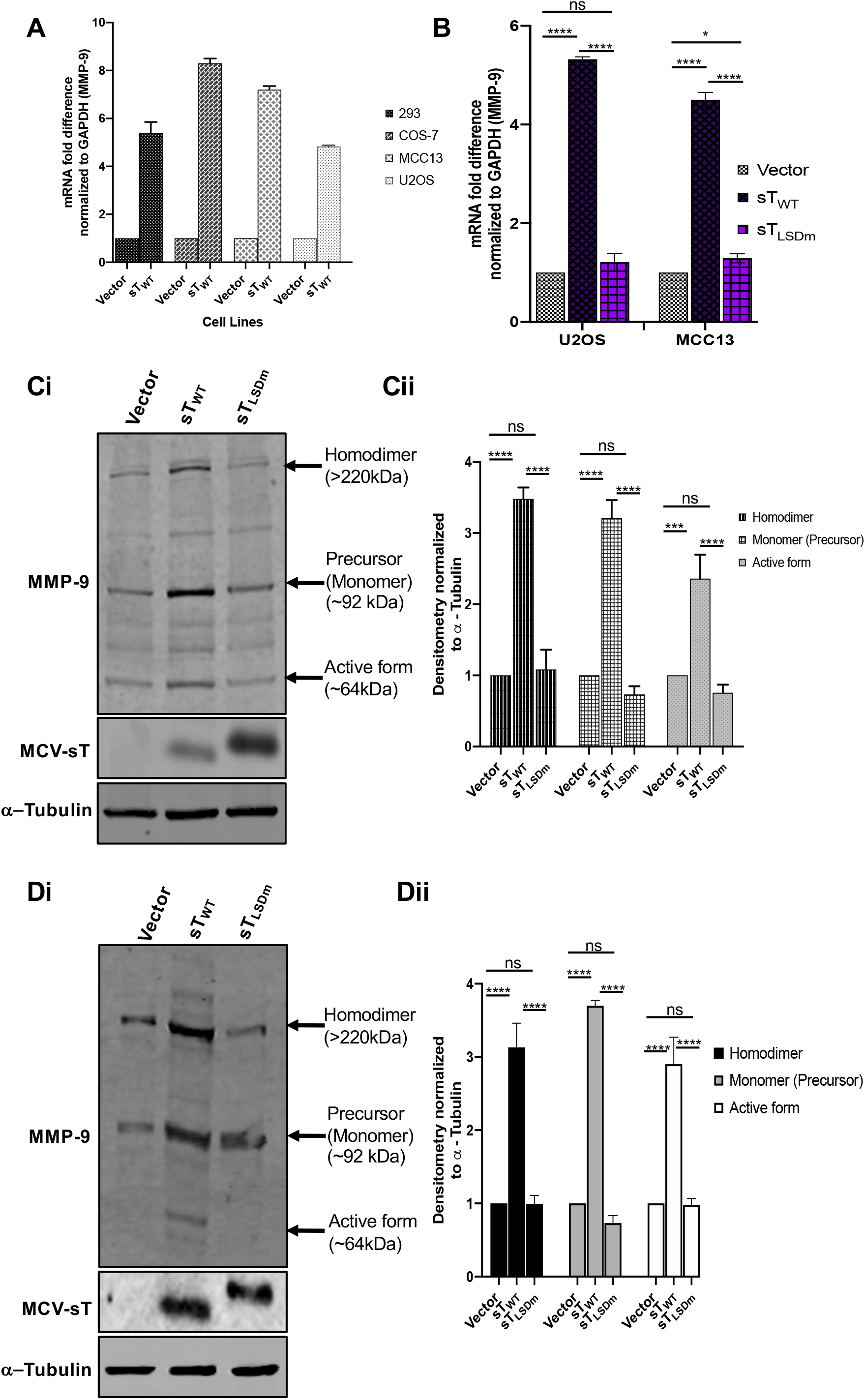
MCV sT activates Matrix metalloproteinase 9 (MMP-9). **(A)** MCV sT expression results in upregulation of MMP-9 mRNA levels. Various cell lines (293, COS-7, MCC13 and U2OS) were transfected with either empty vector or MCV sT_WT_ expressing plasmids to measure MMP-9 mRNA levels. After 48 h, total RNA was isolated and analyzed by RT-qPCR. **(B)** MCV sT upregulates MMP-9 transcription through the LSD. U2OS and MCC13 cells transfected with empty vector control, MCV sT_WT_ and MCV sT_LSDm_ expressing plasmids. Transcript levels of MMP-9 were analyzed using the comparative ΔΔCt method. (n = 3). Differences between means (*p* value) were analyzed using a t-test with GraphPad Prism software. **(C)** MCV sT upregulates MMP-9 protein expression through the LSD. U2OS cells were transfected with empty vector, sT_WT_ and sT_LSDm_ expression plasmids. After 48 h, immunoblot analysis was performed to analyze expression of MMP-9, sT and α-tubulin (Ci). Densitometry quantification of immunoblots was carried out using the Image studio software and is shown as a fold change relative to the loading control α-tubulin (Cii). Data analyzed using three biological replicates per experiment (n = 3). **(D)** MCV sT reproducibly activates MMP-9 expression in MCC13.

We posited that MCV sT targeting of FBW7 plays a role in promoting the migratory potential of MCC by preventing MMP-9 protein turnover. Both transcriptional and post-transcriptional levels of MMP-9 were assessed in the presence of the MCV sT_WT_ or MCV sT_LSDm_ in U2OS and MCC13 cell lines (**Fig. 3B**). Results showed that MCV sT significantly induced the upregulation of MMP-9 transcripts when analyzed by RT-qPCR in both cell lines, which was not observed upon mutation of the MCV sT LSD (**Fig. 3B**). Additionally, we performed immunoblot analysis to evaluate the effect of sT on MMP-9 protein levels (**Fig. 3C and 3D**). Studies have shown MMP-9 exists in several forms; a monomeric pro-(~92 kDa), a disulfide-bonded homodimeric (~220 kDa) and multiple active forms (~67 and 82 kDa) (36). The active and dimeric forms of MMP-9 play a role in the invasive and migratory phenotypes of cancer cells (37, 38). MCV sT_WT_ expression induced the upregulation of the dimer, monomer, and active forms of MMP-9. However, mutation of the LSD prevented MCV sT-mediated upregulation of MMP-9 as expression levels remained comparable with the control, suggesting that MCV sT-mediated upregulation of MMP-9 is LSD-dependent (**Fig. 3**).

### MMP-9 inhibition impedes MCV sT-induced cell migration

We next sought to determine if MMP-9 inhibition would have an impact on MCV sT induced motile and migratory potentials. The migratory phenotype of U2OS cells transfected with vector control, sT_WT_ and sT_LSDm_ was assessed using a scratch assay in the absence or presence of non-cytotoxic concentrations of MMP9-I and MMP9-II inhibitors (**Fig. 4A and Fig. S2A**). MMP-9 inhibition resulted in a significant decrease in the distance traveled by MCV sT expressing cells (**Fig. 4A and 4B**), confirming that the MMP-9 is a critical migratory factor that is regulated by MCV sT. Incubation of both inhibitors showed a slight decrease in the motility of vector control cells, implying that any changes observed in migratory rates of MCV sT expression cells are not due to changes in cell viability or cytotoxicity. Both inhibitors showed a minor impact on the motility of sT_LSDm_ expressing cells, comparable to vector control cells.

**FIGURE 4.**
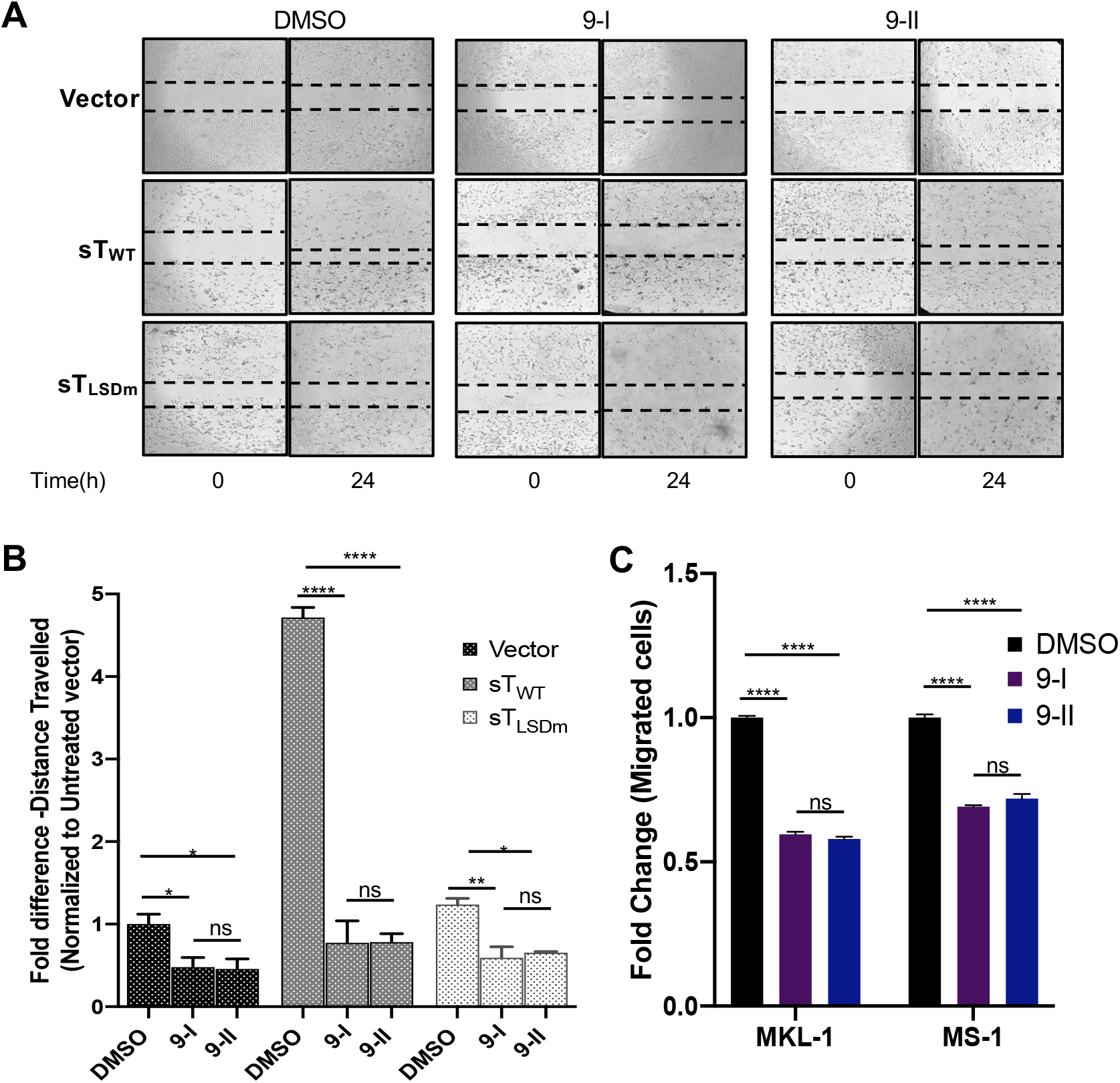
MMP-9 inhibition impedes MCV sT-induced cell migration. **(A)** MCV sT promotes MMP-9-induced cell migration. Scratch assay. Poly-L-lysine-coated 6-well plates were seeded with U2OS cells and incubated with specific MMP-9 inhibitors at predetermined concentrations. Cells were transfected with either a vector control, sT_WT_ and sT_LSDm_ plasmids. After 48 h, a scratch was created and migration of cells toward the scratch was observed over a 24 h period. The size of the wound was measured at 0 and 24 h and presented as the fold change in **(B)**. Scratch assays were performed in triplicate. **(C)** MMP-9 is required for MCC migration. MCV positive MCC cell lines, MKL-1 and MS-1, were incubated with DMSO or the MMP-9 inhibitors 9-I (0.2 μM) and 9-II (2 μM); 9-I (0.1 μM) and 9-II (1 μM), respectively. Cells were then transferred into transwell inserts and allowed to migrate for 48 h. Migrated cells were measured using cell counting kit-8 (CCK-8). Data analyzed using three biological replicates per experiment, n = 3. Differences between means (*p* value) were analyzed using a t-test with GraphPad Prism software.

### MMP-9 is essential for cell motility and migration in MCC

To demonstrate that MMP-9 is vital for cell motility and migration in metastatic MCC, a transwell migration assay was performed using MCV-positive MCC cell lines. This assay quantified the migration ability of MCC cells towards a chemoattractant across a permeable chamber. MCV-positive MCC cell lines, MKL-1 and MS-1 (**Fig. 4C)**, were incubated in the absence or presence of the MMP9-I and MMP9-II inhibitors at non-toxic concentrations (**Fig. S2B**). After treatment, cells were allowed to migrate for 48 h and the total numbers of migrated cells were measured by cell counting Kit-8 assay. Results showed the migration of MCV-positive MCC cell lines was significantly reduced (~40 to 50%) upon incubation of both MMP-9 inhibitors in comparison to the untreated control, suggesting that MMP-9 expression contributes to the migratory capacity of MCV-positive MCC (**Fig. 4C**). Together, these results indicate that MMP-9 is required for MCV sT-mediated cell migration enhancement in MCC.

### MCV sT invasive phenotype is LSD-dependent

The invasiveness of epithelial cancers is a multi-step process and a key hallmark involves the degradation of the basement membrane. Type IV collagen is a major component in most basement membranes. Multiple studies have correlated overexpression of MMPs with not only an enhancement of cell migration and metastasis, but also the invasiveness of cancer cells (13, 15). In particular, MMP-9 is a key protease associated with the degradation of ECM components, including type IV collagen and laminin, which in turn facilitates invasion of tumors into the circulatory system and promotes metastasis. To test if enzymatic activation of MMP-9 is regulated by sT LSD, we evaluated the effect of MCV sT on MMP-9 substrate collagen IV, by immunofluorescence staining in U2OS cells. Our results demonstrate that collagen IV expression in MCV sT_WT_ expressing cells is significantly reduced in comparison to vector control cells, potentially due to MMP-9 activation induced by MCV sT. In contrast, MCV sT_LSDm_ expressing cells did not show a decrease in collagen IV expression (**Fig. 5A, Fig. S3**). Moreover, our regression analysis revealed that collagen IV expression levels are highly correlated with MCV sT_WT_ or sT_LSDm_ expression levels (**Fig. 5B**). To further validate the effect of MCV sT LSD on collagen IV degradation, we performed an invasion assay using collagen pre-coated inserts in U2OS and MCC13 cells. sT_WT_ induced 4 to 5-fold increases in collagen invasion compared to either vector control or sT_LSDm_ (**Fig. 5C**), inferring that MCV sT induces not only cell migration, but also cancer cell invasion through the LSD.

**FIGURE 5.**
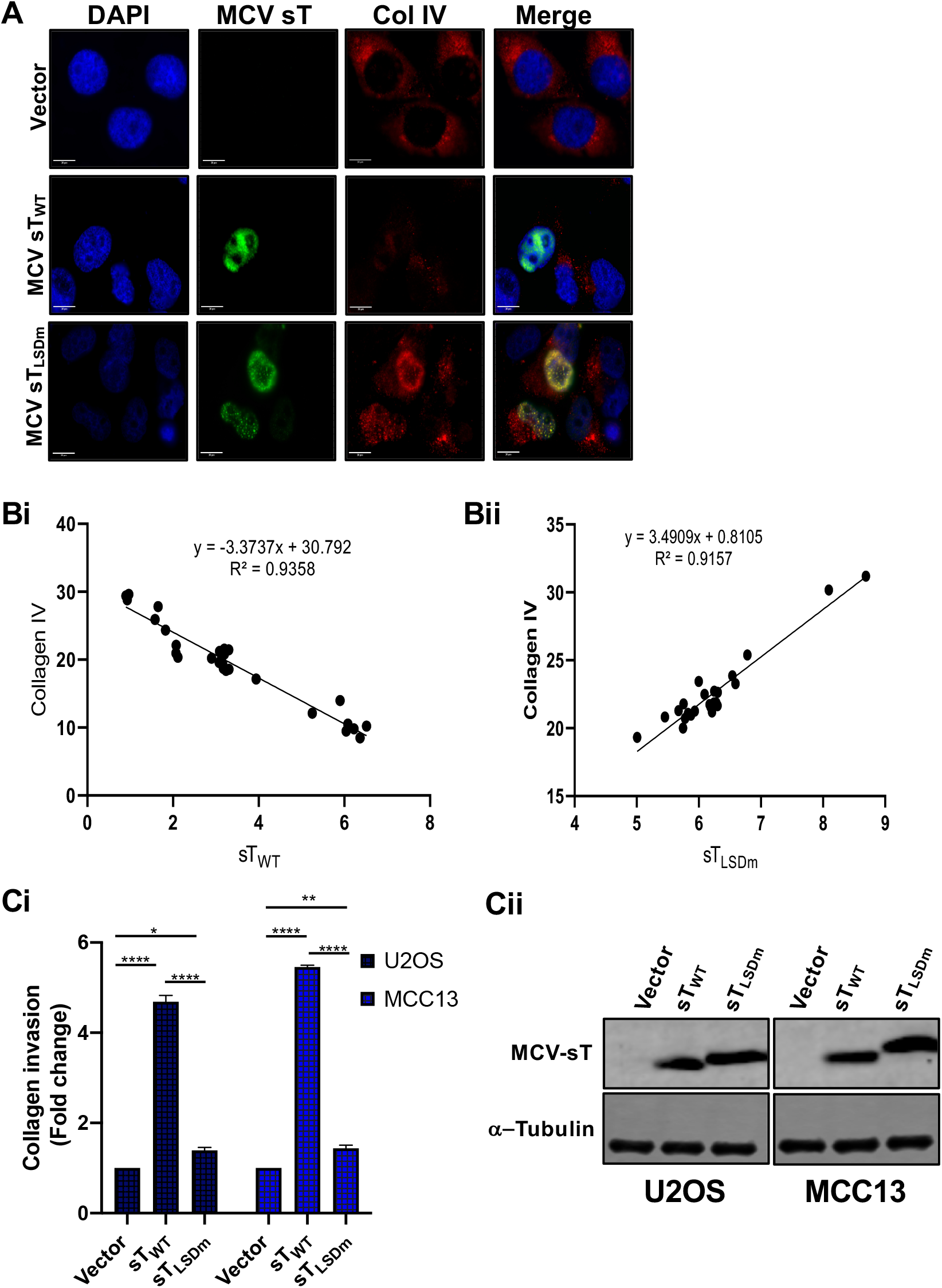
MCV sT LSD induces collagen degradation. **(A)** MCV sT decreases collagen IV expression. U2OS cells were transfected with empty vector control, MCV sT_WT_ and MCV sT_LSDm_ plasmids. Cells were fixed at 48 h post transfection and endogenous collagen IV levels were measured by indirect immunofluorescence using a specific antibody. MCV sT expression was detected with 2T2 antigen antibody. Nuclear counterstain (DAPI-Blue), MCV sT_WT_ and MCV sT_LSDm_ (Green), and collagen IV(Red). **(B)** sT regulates collagen IV expression through the LSD. Mean Florescence intensity of collagen IV in wild type (Bi) and LSD mutant sT-expressing cells (Bii) was analyzed using Fiji Image J software. The calculated values were plotted for regression analysis using Prism software. **(C)** sT induces cell invasion through the LSD. (Ci) Collagen invasion assay. Serum starved sT-expressing U2OS cell were seeded on the precoated collagen inserts and incubated for 72 h, then labeled with a cell staining solution for 20 min. Upon extraction of the cell staining solution, absorbance at OD560 was measured. Data analyzed using three replicates per experiment; the experiments were performed two times. The results were reproducible and differences between means (*p* value) were analyzed using a t-test with GraphPad Prism software. (Cii) Expression of sT. Protein expression levels of wild type and mutant MCV sT were detected by immunoblot analysis to validate successful transfection. Quantitative infrared fluorescence immunoblotting was performed using a 2T2 antibody for sT antigens and α-Tubulin as an equal loading control.

### MCV sT activates expression of EMT regulator, Snail

A positive regulatory loop has been identified between MMP-9 and Snail. siRNA mediated inhibition of MMP-9 significantly reduces expression of Snail, and conversely, knockdown of Snail, a transcription factor of MMP-9, suppresses expression of MMP-9 (39). FBW7 abrogation of Snail protein also inhibits MMP-9 expression (40). Interestingly, MCV sT induced Snail expression in our initial transcriptional analysis (**Fig. 1B**). Since both MMP-9 and Snail are vital mediators of EMT, we assessed the effect of sT LSD on transcriptional and protein levels of Snail in U2OS cells. RT-qPCR results showed that MCV sT_WT_ upregulated mRNA levels of Snail, while this transcriptional change was not observed upon mutation of the LSD (**Fig. 6A)**. Similar to RT-qPCR data, our results demonstrated a significant increase in Snail protein levels upon MCV sT expression in an LSD-dependent manner (**Fig. 6B)**.

**FIGURE 6.**
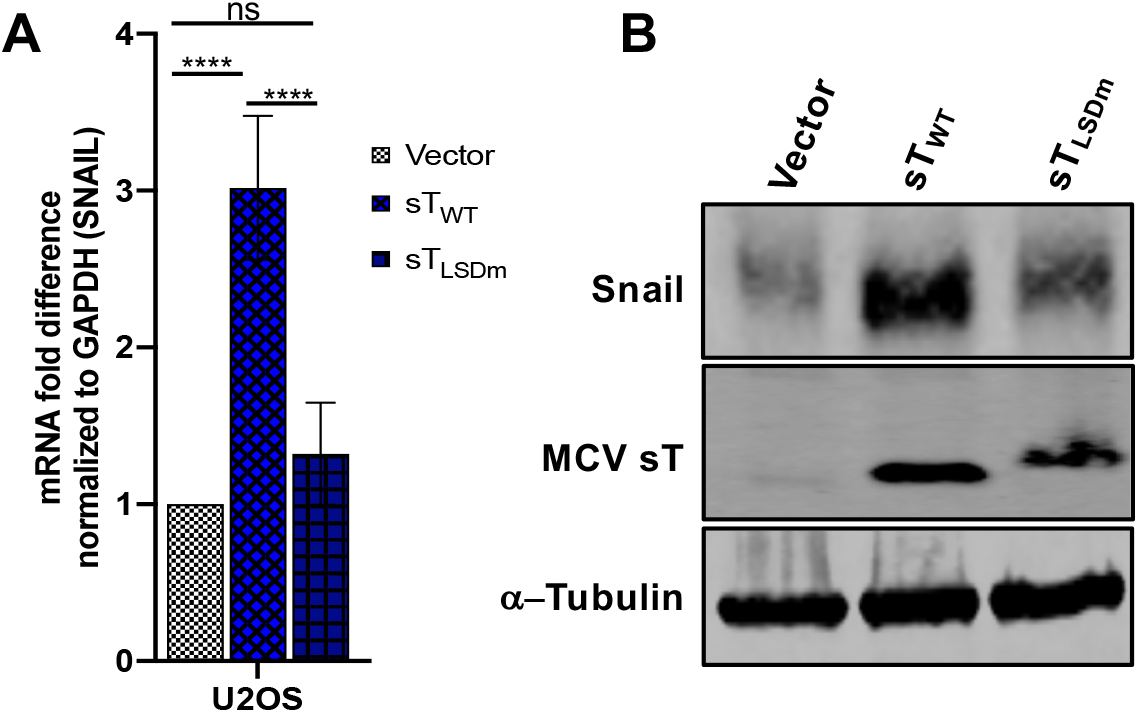
MCV sT LSD induces Snail expression. **(A)** MCV sT activates the transcription of Snail in an LSD-dependent manner. U2OS cells were transfected with empty vector control, sT_WT_ and sT_LSDm_ expressing plasmids. Cellular RNA was extracted at 48 h post transfection and transcript levels were analyzed using the comparative ΔΔCt method (n = 3). **(B)** Snail protein expression is induced by MCV sT through the LSD. Immunoblot analysis was performed on the cellular lysates and analyzed using Snail specific antibody. α-Tubulin was used as a measure of equal loading and the 2T2 antibody was used to confirm MCV sT wild type and mutant expression.

## DISCUSSION

Metastasis is the endpoint of a series of biological processes by which a tumor cell detaches from the primary tumor and disseminates to a distant site through the circulatory system and establishes a secondary tumor (41). Oncogenic viruses often modulate the EMT axis via regulating E-cadherin repression (42–44), fibroblast growth factor (FGF) ligand modulation (44, 45), cadherin switching (46, 47), induction of transcription factors such as TWIST (48, 49) and Snail (42, 50), and MMP-9 upregulation (17–19). These cellular targets can regulate cancer cell migration and invasion; therefore, they could be exploited for therapeutic strategies in virus-induced metastatic cancers.

Multiple F box proteins can function as tumor suppressors by negatively regulating oncoproteins, and various studies have focused on elucidating this mechanism in tumorigenesis and EMT progression (34). In this report, we show that MCV sT LSD inhibition of FBW7 promotes the upregulation of MMP-9 and contributes to MCV sT-mediated cell migration and invasion. The mechanism by which FBW7 regulates MMP-9 expression is currently unclear, although many studies have shown MMP-9 expression is directly and indirectly regulated by FBW7 substrates such as XBP1, Notch1, and Snail (12, 40, 51). Snail is known to be a substrate of both FBW7 (40) and β-TrCP (52), another major SCF E3 ligase that MCV sT targets through the LSD (7). As previously shown, Snail gene expression induces the loss of epithelial markers and the gain of mesenchymal markers, as well as promoting changes in cell motility and invasive properties (53). Our study initially focused on an MMP-9 specific metastatic progression induced by MCV sT due to limited availability of Snail inhibitors. However, the distressed proteome balance in EMT molecules induced by MCV sT might be triggered by Snail activation through MCV sT targeting multiple E3 ligases, which requires further investigation.

While the detailed regulatory mechanisms and specificity of sT function in transcription modulation remain unclear, studies have shown that MCV sT mediates cellular transcriptome/chromatin remodeling (25, 54) which may alter transcriptional activity and gene expression. Consistent with our data, Berrios et al. also reported that MCV sT downregulates extracellular matrix organization and cell adhesion molecules in their transcriptome analysis (25). We have demonstrated that MCV sT specifically activates both mRNA and protein levels of the EMT-related cellular proteins MMP-9 and Snail through the LSD; however, our results do not rule out the possibility that this effect is potentially modulated by multiple mechanisms in MCC. Nonetheless, it is clear that MCV sT LSD plays a critical role in regulating metastasis-initiating capacity in MCC that could be a potential target for therapeutic interventions.

The underlying mechanism for the high propensity of MCC tumors to metastasize is yet to be elucidated. Because of the rare and aggressive nature of metastatic MCC and the lack of standard chemotherapy, there are no prospective studies of outcomes following treatment of distant metastatic MCC. Since recent FDA approvals of Avelumab and Pembrolizumab represent the only approved treatment option for metastatic MCC (55, 56), it is necessary to evaluate the preclinical anticancer activity of efficient and economical chemotherapeutics through retrospective analysis for both MCV-negative and positive MCC patients. MCV-negative MCC tumors patients are more likely to present with advanced disease than patients with virus-positive tumors (66.7% vs. 48.3%) (57). However, targeting the signaling pathways implicated in regulating tumor invasion could be an effective therapeutic protocol for both types of metastatic MCC treatment.

Our study is the first approach to investigate the therapeutic potential of matrix metalloproteinase in MCC. MCV sT specifically activates the EMT-related cellular proteins MMP-9 through the LSD, which we targeted by commercially available inhibitors, and revealed a potential secondary treatment for distant metastatic MCC.

## MATERIALS AND METHODS

### Cells

293, U2OS, and COS-7 cells were maintained in Dulbecco’s modified Eagle’s medium (DMEM) containing 10% fetal bovine serum (FBS) (Seradigm). MCC13, MS-1, and MKL-1 cell lines were maintained in RPMI 1640 medium supplemented with 10% FBS (Seradigm). NIH3T3 cells were maintained in DMEM with 10% bovine calf serum (Seradigm).

### Plasmids, transfection and transduction

Plasmids for vector control, codon-optimized cDNA constructs for sT_WT_ and sT_LSDm_ have been previously described (5). HA-Fbw7 and Flag-cMyc (32) plasmids were kindly provided by Dr. Nakayama (Kyushu University, Japan). FBW7ΔDF(d231-324) was generated by overlapping PCR using primers listed in Table S2. For sT protein expression, cells were transfected using Lipofectamine 3000 (Invitrogen) or jetOPTIMUS (Polyplus Transfection) according to the manufacturer’s protocol. For lentiviral transduction, codon-optimized cDNAs for MCV sT_WT_, MCV sT_LSDm_, (5) and H-RasV12 were inserted into pLVX empty vector. Plasmids used for this study were listed in **Table S1 (Supplemental material)**. For lentivirus production, 293FT (Invitrogen) cells were used for induction according to the manufacturer’s instructions. Cells were selected with puromycin (3 μg/ml) after infection for one week.

### Reverse transcription-quantitative polymerase chain reaction (RT-qPCR)

RNA was extracted using Monarch Total RNA miniprep kit (New England Biolabs), as per the manufacturer’s instruction. 250 ng of RNA was used as a template in each reaction with iTaq Universal One-Step RT-qPCR Kit (Bio-Rad) or Luna Universal One-Step RT-qPCR Kit (New England Biolabs). Primer sequences used are described in **Table S2**. With GAPDH as an internal control, quantitative analysis was performed using the comparative ΔΔCt method.

### Quantitative immunoblotting (IB) and antibodies

Cells were lysed in IP buffer (50 mM Tris-HCl (pH 8.0), 150 mM NaCl, 1% TritonX-100, 1 mM PMSF, 1 mM benzamidine) and sonicated whole cell lysates were used for direct immunoblotting. Primary antibodies were incubated overnight at 4°C, followed by 1 h secondary antibody incubation at RT. All signals were detected using quantitative Infrared (IR) secondary antibodies (IRDye 800CW goat anti-mouse, 800CW goat anti-rabbit, 680LT goat anti-rabbit IgG, 680LT goat anti-mouse IgG) (LI-COR). Signal intensities were analyzed using a laser-scanning imaging system, Odyssey CLX (LI-COR). Antibodies used for this study are listed in **Table S3**. Protein levels were quantitated and normalized by control, α-Tubulin, or β-Actin, using an Odyssey LI-COR IR imaging system.

### SILAC data analysis

The previously published SILAC-based quantitative proteomic data set analyzing host cell proteome changes upon MCV sT expression (21) and further interrogated using the Database for Annotation, Visualization and Integrated Discovery (DAVID) v6.7 (51). For quantitative analysis, a 2.0-fold cutoff was chosen as a basis for investigating potential proteome changes (50).

### Proximity ligation assay (PLA) Flow cytometry

PLA was performed using a Duolink assay kit (Sigma-Aldrich) according to the manufacturer’s instructions. To evaluate MCV sT and FBW7 interaction, HA-FBW7ΔDF(d231-324) was co-expressed with sT_WT_ or sT_LSDm_. FBW7ΔDF was also co-expressed with c-Myc, a known FBW7 substrate, as a positive control (32). Primary antibodies were utilized at optimized concentrations with HA-Tag (C29F4) (1:500), c-Myc (9E10) (1:500), and 2T2 (1:500) (Millipore). Cells were analyzed by flow cytometry on a 16-color BD LSR Fortessa. The acquired data were analyzed using FlowJo software (Tree Star, Ashland, OR, USA).

### Scratch wound-healing assay

Cells were seeded into the Poly-L-Lysine-coated 6-well plates and transfected with either empty vector or sT_WT_ or sT_LSDm_ plasmids. Because MCV sT promotes serum-independent cell growth (58), a serum starvation condition was not considered for our scratch assay to exclude cell proliferation effect by sT. After 48 h, a scratch was created by scraping the monolayer using a p1000 pipette tip. The migration of cells toward the scratch was observed over a 24 h period, and images were taken every 8 h under a REVOLVE4 fluorescent microscope (Echo Laboratories). Inhibitor-based scratch assays were incubated for 24 h prior to transfection with 0.1 and 1 μM of 9-I and 9-II inhibitors respectively.

### Transwell cell migration assay

Cells grown in DMEM with 10% FBS were trypsinized and resuspended in DMEM. 1L×L10^5^ cells were gently added to the transwell insert (8 μm, Greiner Bio-One). DMEM with 10% FBS was added to the bottom of the lower chamber (24-well plate). The cells were incubated in the culture incubator at 37 °C plus 5% CO2 for the indicated time.

The cells migrated from the insert to the well through the filter. The filter was fixed with 4% paraformaldehyde in PBS for 10 min, then stained with 1% Crystal Violet in 2% ethanol for 20 min for NIH3T3 cells and MCC cells were counted using a Cell Counting Kit-8 (CCK-8) (Sigma-Aldrich). The stained cells on the lower side were counted under a microscope from 5 different randomly selected views. All conditions were the same for assays performed in triplicate.

### Immunofluorescence

U2OS cells grown on glass coverslips were transfected with empty vector or sT wild type or LSD mutant expression constructs. After 48 h, cells were fixed in 1:1 methanol/acetone at −20°C, permeabilized, and blocked in PBS with 5% BSA and 0.3 M glycine for 1 h. Cells were labeled with the appropriate primary antibodies and then incubated with the appropriate Alexa Fluor-conjugated secondary antibody. Cells were analyzed with a REVOLVE4 fluorescent microscope (Echo Laboratories).

### Collagen invasion assay

U2OS and MCC13 cells were transfected with wild type and LSD mutant sT constructs for 48 h, followed by overnight serum starvation. 1 ×10^^6^ cells resuspending in serum-free media in each condition were seeded in a 24-well cell invasion plate containing polymerized collagen-coated membrane inserts. The collagen inserts had a pore size of 8 μm (Chemicon QCM Collagen Cell Invasion Assay, ECM551). Complete medium was used as a chemoattractant in the lower chamber and cells were left to incubate for 72 h. Cells/media were carefully aspirated by pipetting any residual suspension in the transwell insert. Inserts were transferred to a clean well and were carefully stained with 400 μL cell staining solution at room temperature for 20 minutes, followed by a gentle wash in deionized water. While slightly damp, unattached cells were removed cautiously by cotton swabs from the collagen inserts and allowed to dry at room temperature for 15 minutes. Dried inserts were transferred to clean wells containing 200 μL of extraction buffer and incubated for 15 minutes at room temperature.

Following the extraction incubation, 100 μL of the extraction solution was pipetted into 96 well plates, and optical density was measured at 560 nm.

### Cell Proliferation Assay

U2OS transfected cells (vector control, MCV sT_WT_ and MCV sT_LSDm_ plasmids) were seeded in 96 well plates (1 × 10^4^ cells/well) 48 h post-transfection. Cell proliferation was monitored using a WST-8 based assay Cell Counting Kit-8 (CCK-8) according to the manufacturer’s protocol. OD values were divided by the OD value of day 0 for normalization.

### Chemical inhibitors

MMP-9 inhibitors-I and II (EMD Millipore) were used at 0.1 to 0.2 μM and 1 to 2 μM, respectively. Cell toxicity was measured using a Cell Counting Kit-8 (CCK-8) (Sigma-Aldrich) according to the manufacturer’s protocol.

### Statistical analysis

Statistical significance between two groups was determined using one- or two-tailed student’s t-tests in GraphPad Prism (GraphPad Software, Inc., La Jolla, CA, USA). The difference was considered significant when p <□0.05 for multiple testing. *, **, *** = p-value < 0.01, 0.005 and 0.001, respectively.

## Acknowledgments

We thank Dr. Patrick S Moore and Dr. Yuan Chang for kind sharing of MCV-related reagents. H.J.K. was supported in part by an Institutional Research Grant, IRG-17-175-04 from the American Cancer Society, and by the Pennsylvania Department of Health Tobacco CURE Funds. N.N. was supported by training grant T32 CA060395 from the National Cancer Institute, National Institutes of Health.

