## Supplemental Material for "Merkel Cell Polyomavirus Small Tumor Antigen Activates Matrix Metallopeptidase-9 Gene Expression for Cell Migration and Invasion"

This PDF file includes:

Figures S1-S3

Tables S1-S3

### Fig S1

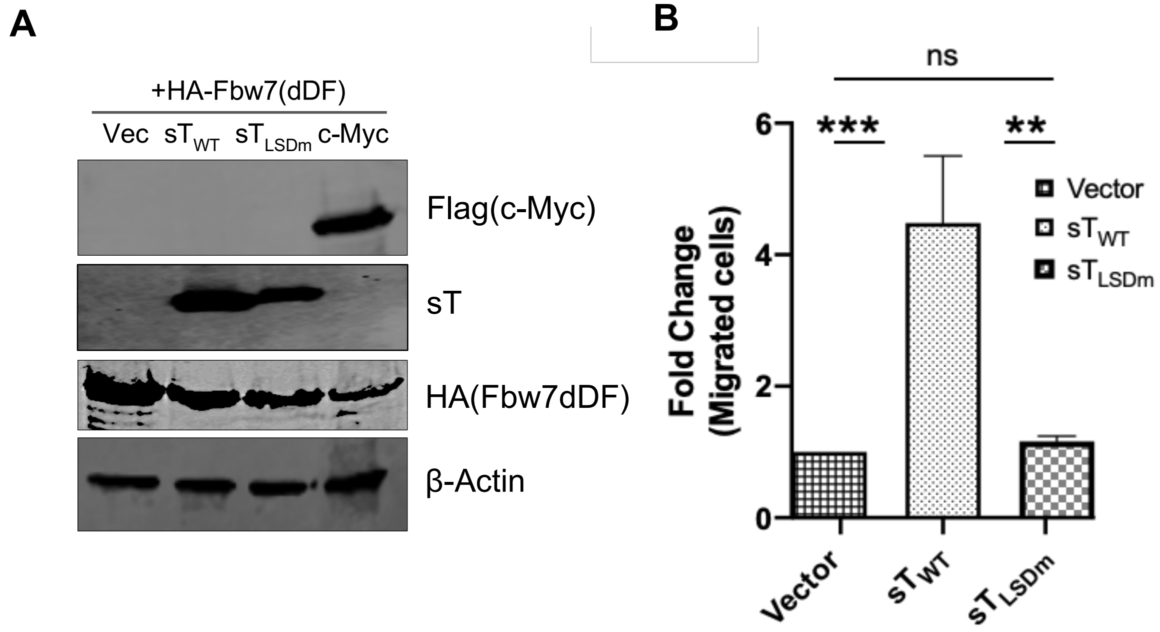


**Fig S1. MCV sT induces cell motility in an LSD-dependent manner. (A)** Protein expression levels of PLA flow cytometry samples used in Fig. 2A. Protein expression was evaluated by immunoblot analysis to validate successful transfection. Quantitative Infrared fluorescence immunoblotting was performed for sT antigen, HA-FBW7, Flag-c-Myc, and β-Actin expression, respectively. **(B)** MCV sT induces MCC cell motility. MCC13 cells stably expressing an empty vector, $sT_{WT}$ and $sT_{LSDm}$ (Fbw7 binding mutant) were trypsinized and $1x10^5$ cells were seeded into transwell inserts and incubated for 3 days. Data analyzed using three replicates per experiment, the experiments were performed two times. The results were reproducible and differences between means (*p* value) were analyzed using a t-test with GraphPad Prism software. Expression of sT protein is shown in Fig.5.

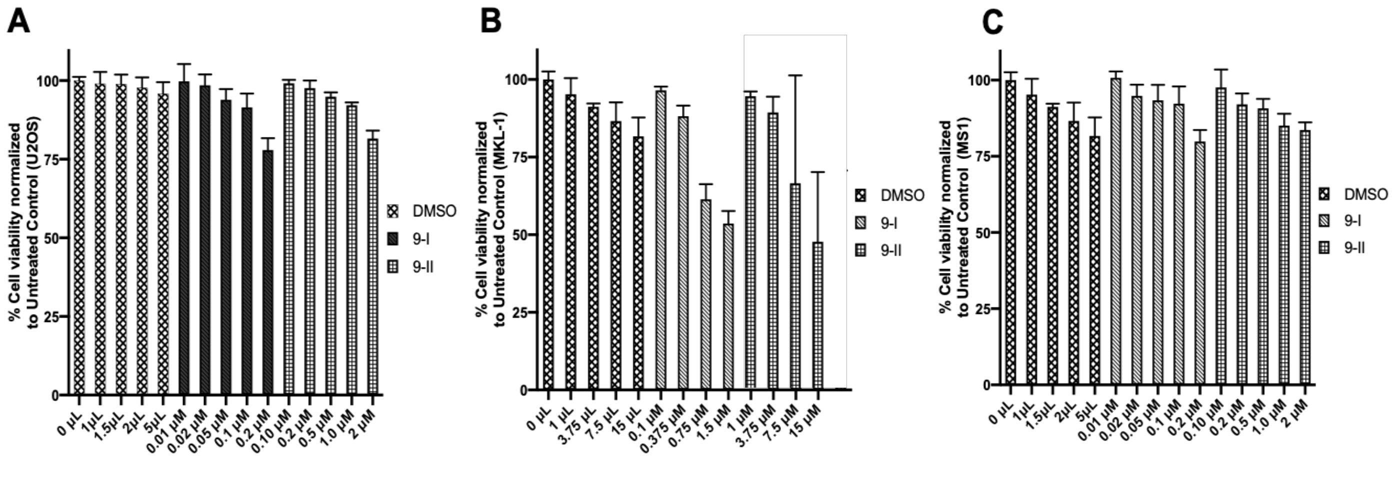


**Fig S2. Cell viability assay for MMP-9 protein inhibitors**. (A) U2OS (B) MKL-1 (C) MS-1 cells (5 x $10^5$ cells) were seeded into 96 well plates and treated with increasing concentrations of (i) MMP-9 homodimer inhibitor (9-I) or MMP-9 hydroxamate active site inhibitor (9-II) for 48 hours. 10 µl of the CCK-8 reagent was added for 4 hours and cell viability was measured at 450nm using a plate reader, Synergy H1.

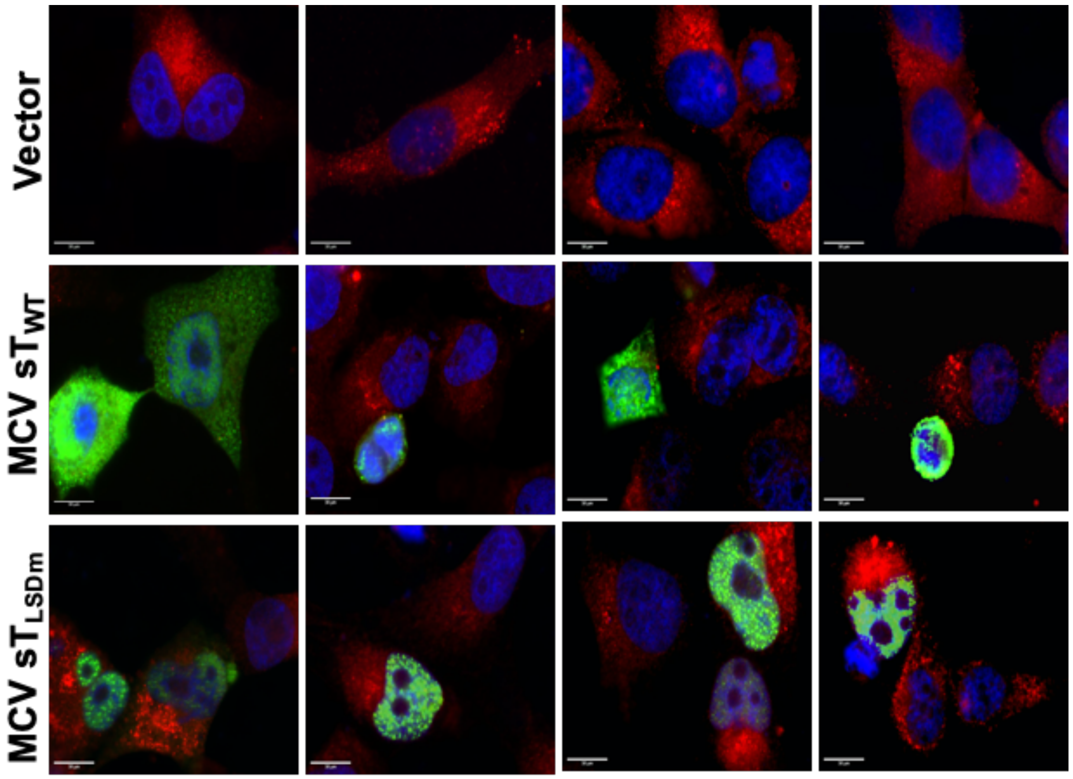


**Fig S3. MCV sT LSD induces a decrease in collagen expression**. U2OS cells were transfected with empty vector control, MCV sT$_{WT}$ and MCV sT$_{LSDm}$ plasmids. Cells were fixed at 48 h post transfection and endogenous Collagen IV levels were measured by indirect immunofluorescence using a specific antibody. MCV sT expression was detected with 2T2 antibody. Merged Images – Nuclear counterstain (DAPI-Blue), MCV sT$_{WT}$/MCV sT$_{LSDm}$ (Green), and Collagen IV (Red).

**Table S1. Plasmids used in this study.**

| Plasmid name | Kwun Lab plasmid # |
|---|---|
| pcDNA.s$T_{WT}$ | 83 |
| pcDNA.s$T_{LSDm}$ | 84 |
| pLVX.H-RasV12 | 8 |
| pLVX EF-MCS Puro | 27 |
| pLVX EF s$T_{WT}$ Puro | 5 |
| pLVX EF s$T_{LSDm}$ Puro | 339 |
| pCI Flag-Myc | 40 |
| pCGN HA-FBW7 (d231-324) WT | 97 |

**Table S2. Primers used in this study.**

| Primers | Sequences |
|---|---|
| MMP-9 Forward | TTT GAG TCC GGT GGA CGA TG |
| MMP-9 Reverse | GCT CCT CAA AGA CCG AGT C |
| GAPDH Forward | CCT CCC GCT TCG CTC TCT |
| GAPDH Reverse | CTG GCG ACG CAA AAG AAG A |
| SNAIL Forward | GCG AGC TGC AGG ACT CTA AT |
| SNAIL Reverse | GGA CAG AGT CCC AGA TGA C |
| AfeI_HRas G12V.F | GCA GCG CTA TGA CGG AAT ATA AGC TGG TG |
| BamHI_HRas G12V.R | CCT GGA TCC TCA GGA GAG CAC ACA CTT GCA |
| Fbw7.d278-324.F | GTG ATA GAA CCC CAG TTT CAA TGC AAA GAA GAG GGG ATT GAT G |
| Fbw7.d278-324.R | GGT TCA TCA ATC CCC TCT TCT TTG CAT TGA AAC TGG GGT TCT ATC ACT TG |

**Table S3: Antibodies used in this study.**

| Name | Supplier | Detection |
|---|---|---|
| Pan Ras (F132) | Santa Cruz Biotechnology | Ras |
| MMP9 (Middle) (ab26132) | Proteintech | MMP-9 |
| Anti-MCPyV T-antigen Antibody (2T2) | Millipore | MCV sT |
| Snail (C15D3) | Cell Signaling Technology | Snail |
| Anti α-Tubulin (12G10) | DSHB | Alpha Tubulin |
| Collagen IV (ab6586) | Abcam | Collagen IV |
| c-Myc (9E10) | DSHB | c-Myc |
| HA-Tag (C29F4) | Cell Signaling Technology | FBW7 construct |
| Anti-Flag (M2) | Sigma | c-Myc |
| β-Actin (13E5) | Cell Signaling Technology | β-Actin |
| IRDye 800CW goat anti-mouse IgG | LI-COR | |
| IRDye 800CW goat anti-rabbit IgG | LI-COR | |
| IRDye 700CW goat anti-mouse IgG | LI-COR | |
| IRDye 700CW goat anti-rabbit IgG | LI-COR | |
| Goat anti-Mouse IgG (H+L) Highly Cross-Adsorbed Sec. Ab, Alexa Fluor 488 (A11029) | ThermoFisher | Secondary immunofluorescent detection of 2T2 |
| Goat anti-Rabbit IgG (H+L) Highly Cross-Adsorbed Sec. Ab, Alexa Fluor 658 (A11036) | ThermoFisher | Secondary immunofluorescent detection of Collagen IV |
